# The persistence of memory: prior memory responses modulate behavior and brain state engagement

**DOI:** 10.1101/2024.04.05.588245

**Authors:** Justin R. Wheelock, Nicole M. Long

## Abstract

Memory brain states may influence how we experience an event. Memory encoding and retrieval constitute neurally dissociable brain states that individuals can selectively engage based on top-down goals. To the extent that memory states linger in time – as suggested by prior behavioral work – memory states may influence not only the current experience, but also subsequent stimuli and judgments. Thus lingering memory states may have broad influences on cognition, yet this account has not been directly tested utilizing neural measures of memory states. Here we address this gap by testing the hypothesis that memory brain states are modulated by memory judgments, and that these brain states persist for several hundred milliseconds. We recorded scalp electroencephalography (EEG) while participants completed a recognition memory task. We used an independently validated multivariate mnemonic state classifier to assess memory state engagement. We replicate prior behavioral findings; however, our neural findings run counter to the predictions made on the basis of the behavioral data. Surprisingly, we find that prior responses modulate current memory state engagement on the basis of response congruency. That is, we find strong engagement of the retrieval state on incongruent trials – when a target is preceded by a correct rejection of a lure and when a lure is preceded by successful recognition of a target. These findings indicate that cortical brain states are influenced by prior judgments and suggest that a non-mnemonic, internal attention state may be recruited to in the face of changing demands in a dynamic environment.

## Introduction

Memory brain states may linger and influence future decisions. For example, imagine reminiscing about graduate school with a close colleague at a conference. If following this conversation you are approached by someone whom you have never met, you may mistakenly find this new colleague familiar. This memory error may be driven by lingering engagement of a memory brain state – specifically, recognizing the known colleague may induce a retrieval state that persists over time and alters your perception of the new colleague. However, the exact dynamics of the interaction between memory brain states and behavior are unclear, leaving open the possibility that shifts in attentional demands may underlie the lingering influence across decisions. The aim of this study was to establish the influence of prior memory judgments on ongoing neural dynamics and decisions.

Memory encoding and memory retrieval states rely on distinct neural substrates and influence how stimuli are processed^1–3^. Rodent electrophysiological work and theoretical models have shown that encoding and retrieval states recruit distinct hippocampal configurations^1, 2^ and both scalp electroencephalography (EEG) and functional magnetic resonance imaging (fMRI) work have shown that multivariate pattern analysis (MVPA) can be used to distinguish between memory encoding and memory retrieval states^4, 5^. Stimuli are selectively represented in either visual or parietal regions depending on which memory state, encoding or retrieval, is engaged^5^ and engaging the retrieval state can impair subsequent memory^4^, ostensibly by preventing engagement of the encoding state. Given that these memory states can be engaged in the absence of episodic memory demands^3^ and automatically in the presence of temporal contextual overlap^6^, there is a strong potential for these states to influence cognition. Yet, how and whether these states persist over time and influence not only the current stimulus or judgment but subsequent stimuli remains a critical open question.

Behavioral evidence suggests that memory states may linger on the order of milliseconds to influence memory judgments^7, 8^. Specifically, when participants make an “old” judgment to a previously studied item (target), if a similar lure – a stimulus that is categorically related to a previously studied item – is presented within 500ms of that judgment, the likelihood that the participant will respond “old” to the similar lure increases, relative to if the participant had made a “new” judgment to a novel item on the prior trial. The interpretation is that the “old” judgment to the first probe induces a hippocampally-mediated retrieval state which lingers such that when the second probe is presented, the persistent retrieval state leads to an “old” judgment. Although hippocampal memory states are governed by theta (4Hz) oscillations, these behavioral effects are thought to be the result of a slower, cholinergic modulatory signal that operates on the order of seconds^8–10^. When participants are explicitly instructed to engage the retrieval state, a putative correlate of the default mode network shows sustained engagement across the multi-second stimulus interval^11^, suggesting that cortical memory states may persist along these time scales. However, whether test-phase memory judgments induce a lingering memory state that influences subsequent judgments is unclear.

Our hypothesis is that memory brain states are modulated by memory judgments and that memory brain states persist for several hundred milliseconds, impacting subsequent memory judgments. To test this hypothesis, we conducted a human scalp EEG study in which participants completed a recognition memory task (Figure 1). Our primary manipulation was the interstimulus interval (ISI) between test-phase probes. Using a cross-study decoding approach with independently collected mnemonic state data^3^, we assessed memory state engagement during memory judgments. We expected to replicate prior findings that memory judgments are influenced by the prior response selectively for short ISIs. We predicted that memory brain state engagement would be driven by both the current and prior judgment, whereby “old” judgments would induce a lingering retrieval state and “new” judgments would induce a lingering encoding state.

**Figure 1.**
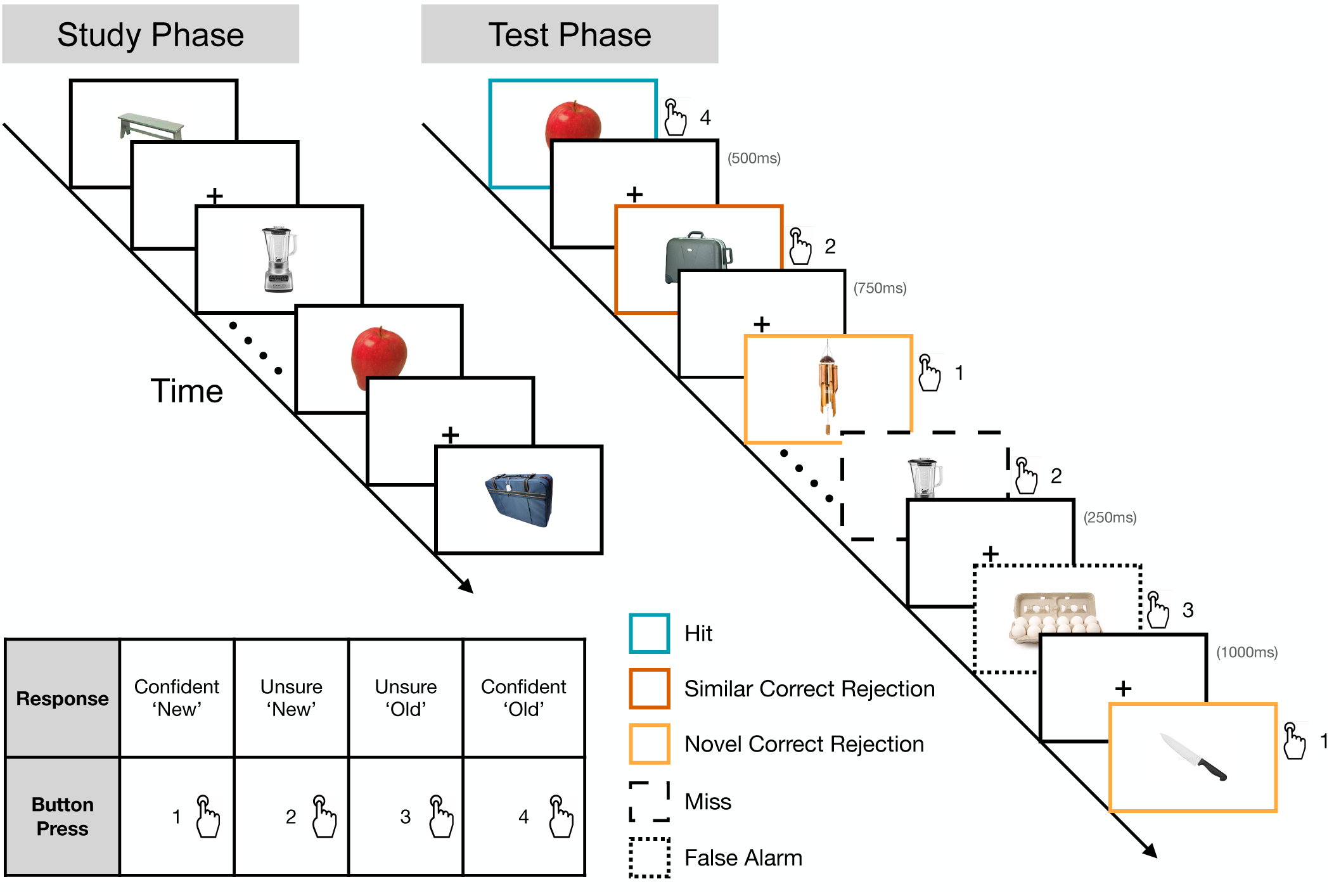
Experimental Design. During the study phase, participants studied 16 lists of 18 images of common objects in anticipation of a later memory test. After completing the study phase, participants completed a recognition test-phase. During each trial, participants were shown either a target (an item that had been presented in the study phase) a similar lure (an item that had not been presented in the study phase but belonged to the same object category as a study-phase item), or novel item (an item that had not been presented in the study phase and was from a unique object category that had not been studied). Participants responded on a scale of 1 to 4, whereby 1 indicated a confident ‘new’ judgment, and 4 indicated a confident ‘old’ judgment. Responses of 2 and 3 were uncertain ‘new’ and ‘old’ judgments, respectively.

## Results

### Prior responses modulate recognition memory accuracy and confidence

Our first goal was to replicate prior work showing that recognition accuracy on the current trial is modulated by prior responses and that this modulation is specific to inter-stimulus intervals (ISIs) of 500ms or faster^7^. To the extent that hits induce a retrieval state, a prior hit should increase the tendency to respond ‘old’ on the current trial, leading to higher hit rates for targets and lower CR rates for lures. Likewise, to the extent that CRs induce an encoding state, a prior CR should increase the tendency to respond ‘new’ on the current trial, leading to lower hit rates for targets and higher CR rates for lures. Thus, we should find a significant interaction between prior response and current probe on recognition accuracy.

To test the influence of prior responses and ISI on the current response, we conducted a 3*×*2*×*2 repeated measures ANOVA (rmANOVA), with current probe (target, similar lure, novel item), prior response (hit, CR), and ISI (fast, slow) as factors and accuracy as the dependent variable (Figure 2A). To maximize the number of trials available for analysis, for the prior response conditions we combine both similar and novel CRs and refer to these trials collectively as “prior CRs.” We report the results in Table 1 and highlight the key findings here. We find a significant main effect of current probe driven by greater accuracy for novel items (M=0.9062, SD=0.0947) compared to both targets (M=0.7758, SD=0.1117) and similar lures (M=0.7758, SD=0.1133; novel vs. target: *t* _26_=5.999, *p<*0.0001, *d* =1.264; novel vs. similar lure: *t* _26_=10.56, *p<*0.0001, *d* =1.254; FDR corrected). We find a significant interaction between current probe and prior response (*F* _2,52_=5.143, *p*=0.0092, 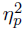=0.1651). We do not find a significant three-way interaction between current probe, prior response, and ISI and Bayes Factor analysis revealed that a model which excludes ISI and the three way interaction term is preferred to a model which includes ISI by a factor of 2.1*×*10^4^. Given the preference for the model that excludes ISI, we average over ISI for all subsequent behavioral analyses.

**Figure 2.**
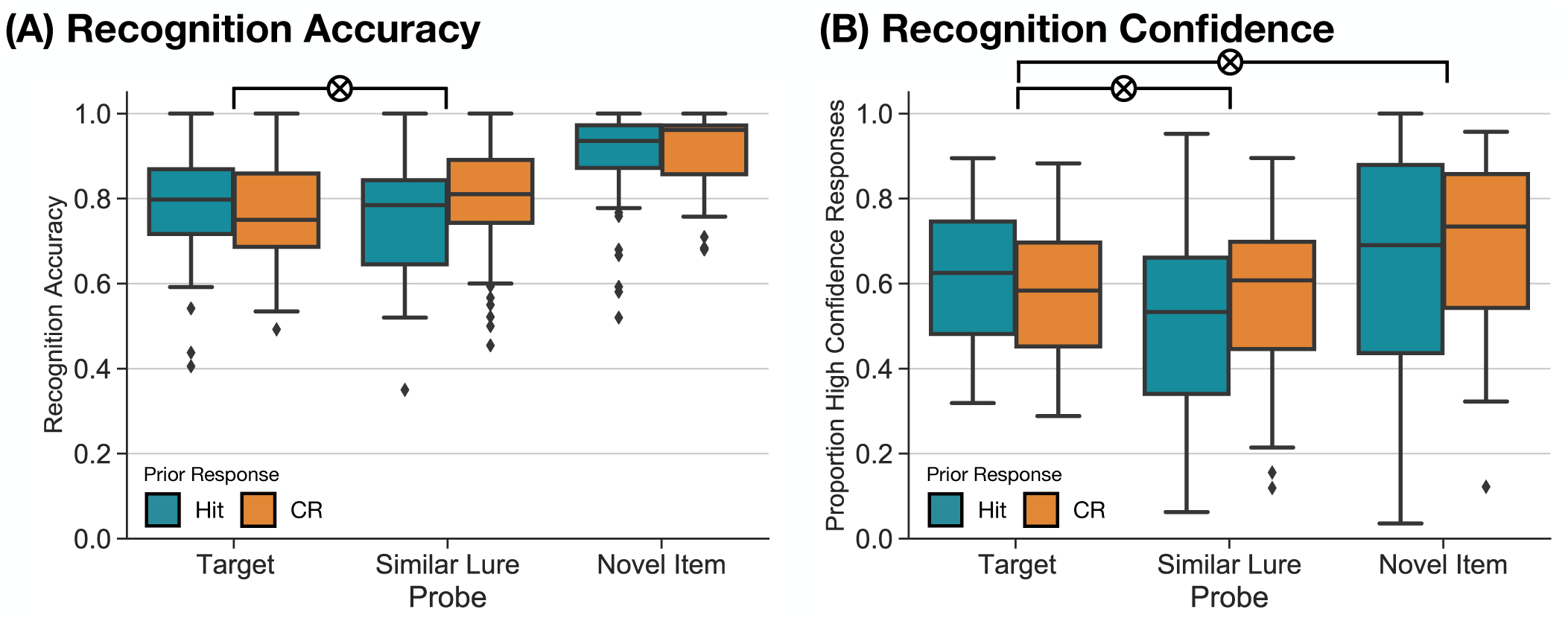
Memory responses impact subsequent judgments. Box-and-whisker plots show median (center line), upper and lower quartiles (box limits), 1.5x interquartile range (whiskers) and outliers (diamonds). **(A)** Recognition accuracy is shown across three probe types (target, similar lure, novel item) based on prior response (prior hit, orange; prior correct rejection, CR, teal). There is a significant interaction between current probe and prior response, driven by the difference between targets and similar lures (*p*=0.009), whereby targets are more accurately identified following hits and similar lures are more accurately identified following CRs. **(B)** The proportion of high confidence responses is shown as a function of current probe and prior response. There is a significant interaction between current probe and prior response (*p<*0.0001), which is driven both by the difference between hits and similar lures, as well as the difference between hits and novel items.

**Table 1.** Influence of prior response on current memory accuracy.

| Effect | df | *F* | *p* | $\eta_p^2$ |
| --- | --- | --- | --- | --- |
| Main effect of current probe | 2,52 | **29.28** | **<0.0001** | **0.53** |
| Main effect of prior response | 1,26 | 3.891 | 0.0593 | 0.13 |
| Main effect of ISI | 1,26 | 0.0605 | 0.8076 | 0.002 |
| Interaction of current probe × prior response | 2,52 | **5.143** | **0.0092** | **0.1651** |
| Interaction of current probe × ISI | 2,52 | 0.5489 | 0.5809 | 0.0207 |
| Interaction of prior response × ISI | 1,26 | 0.1486 | 0.7030 | 0.0057 |
| Interaction of current probe × prior response × ISI | 2,52 | 0.3097 | 0.7350 | 0.0118 |

Bold values indicate tests for which $p < 0.05$

The interaction between prior response and current probe indicates that how participants respond on the current trial is differentially influenced by their response on the prior trial. To test which of the three probes drive this interaction, we calculated difference scores for each probe and conducted three paired *t* -tests on these difference scores. Specifically, we calculated the difference in recognition accuracy for probes preceded by hits minus probes preceded by CRs, separately for targets, similar lures, and novel items. Thus, positive values indicate better performance when the prior response was a hit and negative values indicate better performance when the prior response was a CR. We find that the interaction between prior response and current probe is driven by a significant difference in difference scores between targets and similar lures (*t* _26_=2.873, *p*=0.008, *d* =0.8855; FDR corrected), whereby targets are more accurately identified when preceded by a hit than a CR (difference score: M=0.0187, SD=0.0551), and similar lures are more accurately identified when preceded by a CR than a hit (difference score: M=-0.0431, SD=0.0819). We do not find a significant difference in difference scores between targets and novel items (*t* _26_=1.87, *p*=0.0728, *d* =0.6159), nor between similar lures and novel items (*t* _26_=-1.507, *p*=0.1439, *d* =0.3483). These results indicate that prior responses selectively modulate accuracy on the current trial.

Given that recognition accuracy is modulated by the prior response, we next tested the extent to which recognition confidence is modulated by prior response. We predicted that participants would commit more high confidence responses to targets preceded by hits and to lures preceded by CRs. To assess the effect of prior response on confidence, we calculated the proportion of high confidence hits and high confidence CRs. We conducted a 3*×*2 rmANOVA (Figure 2B), with current probe (target, similar lure, novel item) and prior response (hit, CR) as factors, and the proportion of high confidence responses as the dependent variable. We find a significant main effect of current probe (*F* _2,52_=9.658, *p*=0.0003, 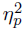=0.2709), whereby participants commit fewer high confidence responses to similar lures (M=0.5178, SD=0.2084), compared to targets (M=0.5992, SD=0.1570) or novel items (M=0.6627, SD=0.2317; target vs. similar lure: *t* _26_=2.294, *p*=0.0301, *d* =0.4411; similar lure vs. novel item: *t* _26_=8.064, *p<*0.0001, *d* =0.6577). We do not find a significant difference in the proportion of high confidence responses to targets vs. novel items (*t* _26_=1.543, *p*=0.135, *d* =0.3211). We find a significant main effect of prior response (*F* _1,26_=6.231, *p*=0.0192, 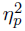 =0.1933), whereby participants commit more high confidence responses following CRs (M=0.5787, SD=0.2359) compared to hits (M=0.5787, SD=0.2359). We find a significant interaction between current probe and prior response (*F* _2,52_=10.72, *p*=0.0001, 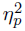=0.292).

As with the accuracy analysis, we find that prior response interacts with the current probe to modulate confidence. To identify which probes drive this interaction, we again calculated difference scores for each probe whereby positive values indicate a greater proportion of high confidence responses following hits, and negative values indicate a greater proportion of high confidence responses following CRs. We find that the interaction between current probe and prior response is driven by both a difference in difference scores between targets and similar lures (*t* _26_=4.613, *p*=0.0001, *d* =1.364) and difference scores between targets and novel items (*t* _26_=3.402, *p*=0.0022, *d* =1.039; FDR corrected). Specifically, participants committed more high confidence responses to targets preceded by a hit compared to targets preceded by a CR (difference score: M=0.0459, SD=0.0655), whereas for both similar lures and novel items, participants committed more high confidence responses when the trial was preceded by a CR compared to when the trial was preceded by a hit (similar difference score: M=-0.0754, SD=0.1074; novel difference score: M=-0.0541, SD=0.1193). We do not find a significant difference in difference scores between similar lures and novel items (*t* _26_=-0.7574, *p*=0.4556, *d* =0.1877). These results indicate that confidence, like accuracy, is modulated by prior responses.

### Current judgments modulate memory states

Our central goal was to test the extent to which memory judgments modulate and are modulated by memory brain states. Based on prior work, we expected hits to induce a retrieval state and CRs to induce an encoding state. Furthermore, we expected these judgment-induced brain states to persist in time following the response and modulate both the behavioral and neural responses on the subsequent trial. That is, the current trial should exhibit more retrieval evidence when preceded by a hit compared to when the current trial is preceded by a CR. To measure memory state engagement, we trained a multivariate pattern classifier to distinguish encoding and retrieval states based on an independently collected dataset^3, 11^. We tested this classifier on test-phase data in the current recognition study.

We first tested the extent to which hits induce a retrieval state and CRs induce an encoding state. We measured response-locked memory state evidence as a function of the current response (Figure 3A). We conducted a 3*×*4*×*15 rmANOVA with current response (hit, novel CR, similar CR), ISI (250ms, 500ms, 750ms, and 1000ms), and time interval (50ms intervals from 500ms pre-response to 250ms post-response) as factors, and memory state evidence as the dependent variable. We restricted post-response time to 250ms to account for trials with the shortest ISI. We find a significant main effect of current response (*F* _2,52_=4.742, *p*=0.0128, 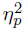=0.1542), driven by greater retrieval evidence for similar CRs (M=0.0444, SD=0.0491) compared to hits (M=0.0205, SD=0.0429; *t* _26_=2.857, *p*=0.0083, *d* =0.5186; FDR corrected). We do not find a significant difference in retrieval state evidence between either hits and novel CRs (M=0.0368, SD=0.0440; *t* _26_=-1.848, *p*=0.076, *d* =0.376) or between similar CRs and novel CRs (*t* _26_=1.187, *p*=0.246, *d* =0.163). We do not find a significant main effect of ISI (*F* _1,26_ = 0.0003, *p*=0.9857, 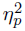*<*0.0001). We find a significant main effect of time interval (*F* _14,364_=7.786, *p<*0.0001, 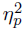=0.2305), whereby retrieval evidence is greater preceding compared to following the response. We find a significant interaction between current response and time interval (*F* _28,728_=1.671, *p*=0.0169, *η*^2^=0.0604). We do not find a significant interaction between current response and ISI (*F* _2,52_=0.2855, *p*=0.7528, 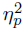=0.0109) or between ISI and time interval (*F* _14,364_=0.8261, *p*=0.6406, 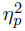=0.0308). We do not find a significant three-way interaction between current response, ISI, and time interval (*F* _28,728_=0.8512, *p*=0.6885, 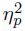=0.0317), and Bayes Factor analysis revealed that a model without ISI and the three way interaction term is preferred to a model with the three way interaction term by a factor of 2.75*×*10^32^. Given the lack of significant ISI effects, we average over ISI for all subsequent neural analyses. Together, these findings directly contradict our *a priori* expectations. Instead of finding greater retrieval evidence for hits compared to CRs, we find greater retrieval evidence for similar CRs compared to hits.

**Figure 3.**
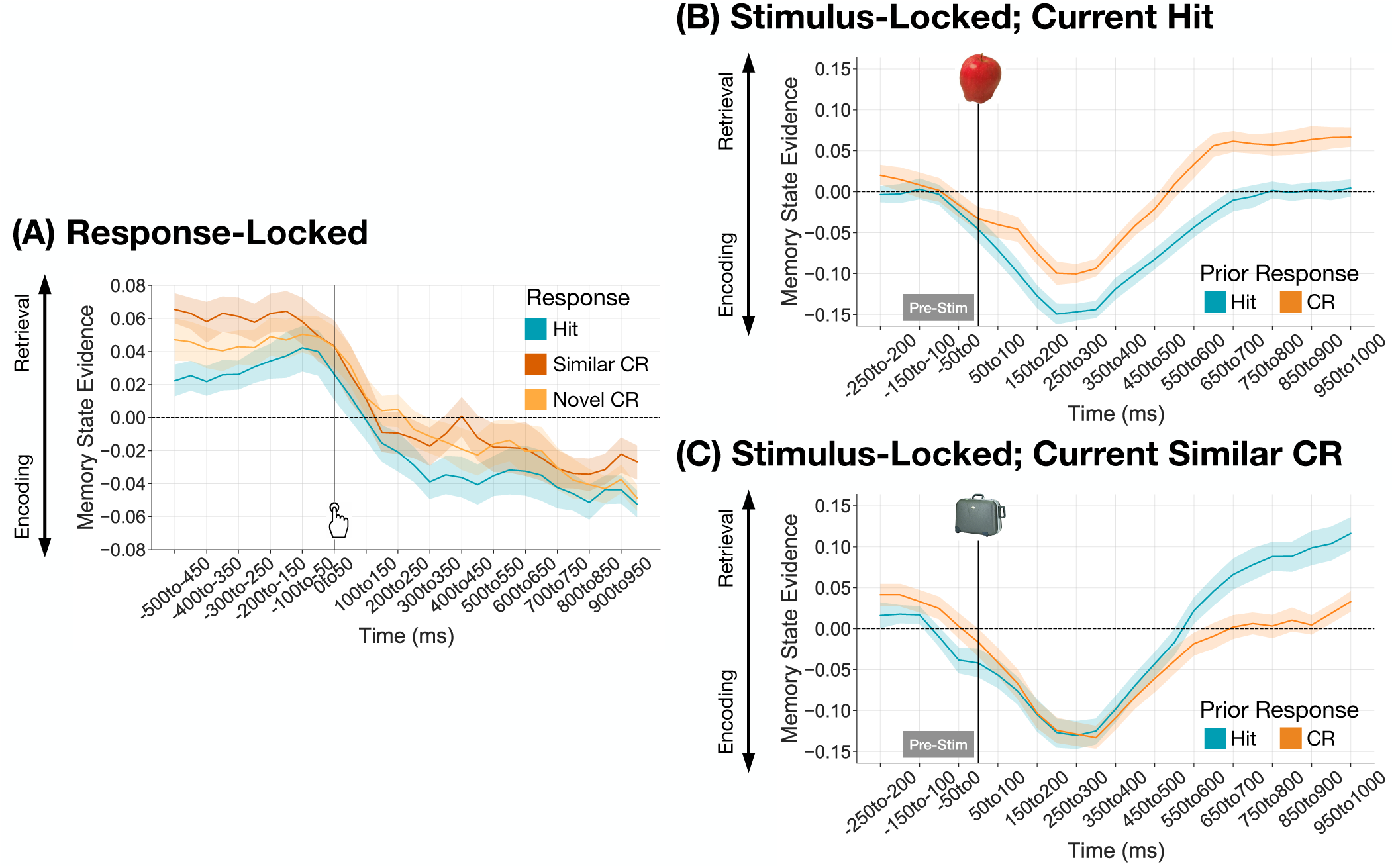
Memory state engagement is modulated by both prior response and current response. Each panel shows memory state evidence over 50 ms time intervals locked to either response onset (A) or stimulus onset (B, C). Shaded areas reflect standard error of the mean. **(A)** Response-locked memory state evidence is shown as a function of response (similar CRs, dark orange; novel CRs, light orange; hits, teal) over thirty time intervals, from 500ms pre-response to 1000ms post-response. There is more retrieval evidence for similar CRs compared to hits (*p*=0.0083). **(B)** Stimulus-locked memory state evidence is shown for current hits as a function of prior response (hit, teal; CR, orange) over twenty-five time interval from 250ms pre-stimulus to 1000ms post-stimulus. There is more retrieval evidence for hits preceded by a CR compared to current hits preceded by a hit (*p<*0.0001). **(C)** Stimulus-locked memory state evidence is shown for current CRs as a function of prior response (hit, teal; CR, orange) over twenty-five time interval from 250ms pre-stimulus to 1000ms post-stimulus. There is more retrieval evidence for current CRs preceded by a hit compared to current CRs preceded by a CR (*p*=0.0035).

**Figure 4.**
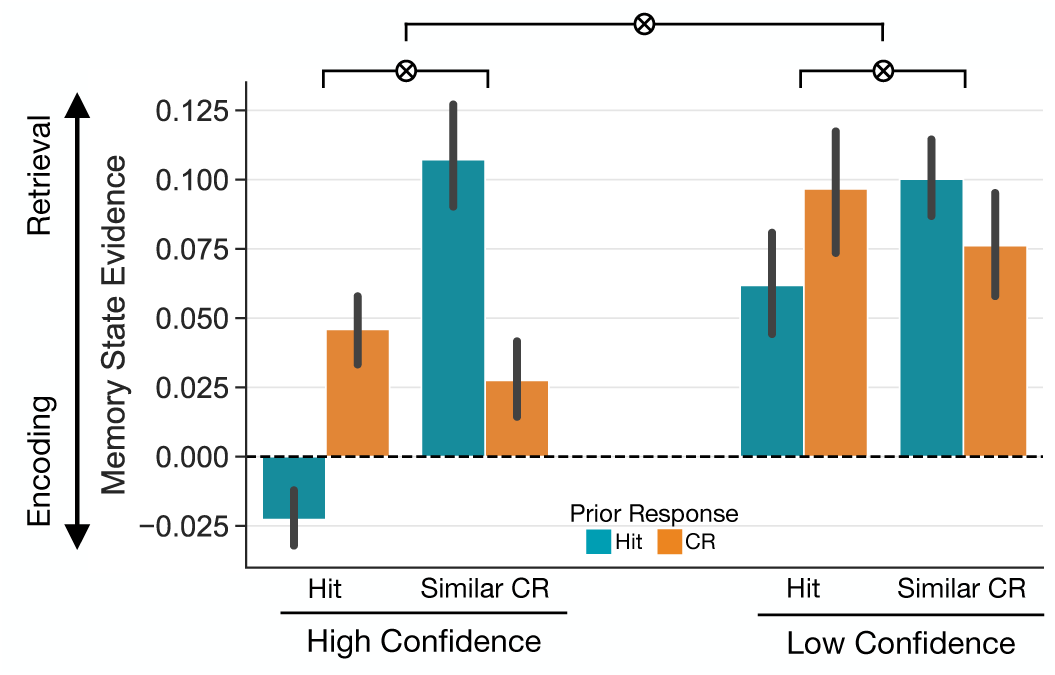
Memory state engagement is modulated by response confidence. Response-locked memory state evidence is shown averaged over the 500ms pre-response interval, as a function of current response (hit or similar CR), prior response (hit, teal; CR, orange), and confidence (high, responses of 1 or 4; low, responses of 2 or 3). There is a significant three-way interaction between current response, prior response, and confidence (*p*=0.0255). There is less retrieval evidence for congruent trials (i.e. hit followed by hit, CR followed by CR) compared to incongruent trials (hit followed by CR, CR followed by hit), and the magnitude of these differences is greater for high confidence trials. Error bars reflect standard error of the mean.

### Limited evidence for lingering memory states

We find that retrieval state evidence decreases immediately following both hits and CRs (Figure 3A) which is counter to the hypothesis that memory states linger following a memory response. However, response-locked data may obscure effects that immediately precede the next trial. Therefore, to directly test the extent to which memory states linger over time, we measured *stimulus-locked* memory state evidence in the 200 ms leading up to stimulus onset separated by the response on the prior trial (Figure 3B,C, “PreStim” interval). We selected 200 ms as the pre-stimulus window as it allows us to directly compare prestimulus memory state evidence without measuring signals during the prior response. With this approach, we should find greater retrieval state evidence for trials preceded by a hit compared to trials preceded by a CR. We conducted a paired *t* -test between pre-stimulus memory state evidence for prior hits and prior CRs. We find that pre-stimulus memory state evidence does not significantly differ between prior hits (M=-0.0024, SD=0.0621) and prior CRs (M=0.0189, SD=0.0685; *t* _26_=1.187, *p*=0.2461, *d* =0.1625). These results thus do not provide evidence to support the hypothesis that memory states linger following a memory judgment.

### Both prior and current responses modulate memory state engagement

Having shown that memory responses modulate memory states – albeit not in the predicted direction – we next sought to test the extent to which both prior and current responses modulate memory states. Although the cortical memory brain states that we measure here may not linger from the prior response into the current trial, the type of response made on the prior trial may still differentially impact current brain state engagement, given the observed behavioral dissociations. That is, although we do not find evidence for selective pre-stimulus memory state engagement, we may still find greater retrieval state engagement for current hits preceded by hits and greater encoding state engagement for current CRs preceded by CRs.

To investigate the modulation of prior response on current memory states, we examined stimulus-locked memory state evidence as a function of the current response (Figure 3B,C). We conducted a 2*×*2*×*20 rmANOVA with factors of prior response (hit, CR), current response (hit, similar CR), and time interval (50ms intervals from stimulus onset to 1000ms post stimulus). We chose to exclusively investigate current hits and similar CRs given that they drive both the behavioral interaction between prior response and current probe and the neural main effect of current response. We chose the interval of 0-1000ms as the highest proportion of test-phase responses across all probes (M=0.76, SD=0.14) occurred after 1000ms and we sought to minimize the contribution of response-related signals in this analysis. We do not find a significant main effect of prior response (*F* _1,26_=3.741, *p*=0.064, 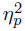=0.1258) or current response (*F* _1,26_=1.191, *p*=0.2852, 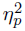=0.0438). We find a significant main effect of time interval (*F* _19,494_=41.17, *p<*0.0001, 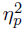=0.6129). We find significant interactions between prior response and current response (*F* _1,26_=35.95, *p<*0.0001, 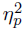=0.5803), prior response and time interval (*F* _19,494_=3.577, *p<*0.0001, 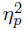=0.1209), and current response and time interval (*F* _14,494_=1.796, *p*=0.0208, 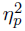=0.0646). Finally, we find a significant three way interaction between prior response, current response, and time interval (*F* _14,494_=7.953, *p<*0.0001, 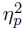=0.2342).

To determine the direction in which both prior and current responses modulate memory states, we conducted separate rmANOVAs for current hits (Figure 3B) and current similar CRs (Figure 3C). Each rmANOVA had factors of prior response (hit, CR) and time interval. For current hits, we find a significant main effect of prior response (*F* _1,26_=42.52, *p<*0.0001, 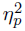=0.6206), whereby retrieval evidence is greater for hits preceded by a CR (M=-0.0042, SD=0.0942) compared to hits preceded by a hit (M=-0.0612, SD=0.0836). We find a significant main effect of time interval (*F* _19,494_=33.94, *p<*0.0001, *η*^2^=0.5662), and a significant interaction between prior response and time interval (*F* _19.494_=1.912, *p*=0.0116, 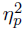=0.0685). For current similar CRs, we find a significant main effect of prior response (*F* _1.26_=10.30, *p*=0.0035, 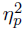=0.2837), whereby retrieval evidence is greater for CRs preceded by a hit (M=-0.0090, SD=0.1255) compared to CRs preceded by a CR (M=-0.0427, SD=0.0927). We find a significant main effect of time interval (*F* _19,494_=33.67, *p<*0.0001, 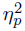=0.5643), and a significant interaction between prior response and time interval (*F* _14,494_=8.694, *p<*0.0001, 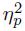=0.2506). Thus, we find that the prior response modulates memory state engagement for both current hits and current similar CRs, yet not entirely in the predicted direction. Retrieval state engagement is greater for CRs preceded by a hit vs. a CR, which is consistent with the expectation that hits induce a retrieval state. However, this explanation cannot account for the pattern that we observe for hits whereby retrieval evidence is greater for hits preceded by CRs. Instead, these findings suggest that *congruency* across consecutive trials may drive memory state engagement.

We have shown that the influence of the prior response on the current memory state depends on the current response. Specifically, there is more stimulus-locked retrieval state evidence when the current and prior response do not match, or, conversely, there is less stimulus-locked retrieval state evidence when the current and prior response match. That is, retrieval state evidence is greater during incongruent compared to congruent trials. To directly test this account, we re-coded trials as congruent (hits preceded by hits and CRs preceded by CRs) or incongruent (hits preceded by CRs and CRs preceded by hits) and averaged retrieval state evidence across the 1000 ms stimulus interval. We find that retrieval state evidence is significantly greater for incongruent (M=-0.0066, SD=0.038) compared to congruent (M=-0.0519, SD=0.0324) trials (*t* _26_=-5.996, *p<*0.0001, *d* =1.283). Together, these results point to a potential influence of switching on memory state engagement. When the probe switches between two consecutive test trials, the retrieval state is more strongly engaged compared to when the probe stays the same.

### Memory state dissociations reveal a mnemonic congruency effect

We have found that the retrieval state, rather than being selectively induced by hits relative to CRs, is specifically induced when consecutive trials are incongruent. That is, hits preceded by CRs and CRs preceded by hits recruit the retrieval state to a greater extent than hits preceded by hits and CRs preceded by CRs. Such an effect could be driven by switches in non-mnemonic task and/or response demands. Although the experimenter-imposed task remained the same throughout this study (recognize old items), evaluating targets and lures may require distinct processes whereby evaluating targets recruits cognitive control and default-mode networks and evaluating lures recruits encoding and priming mechanisms^12^. Thus, the change from either a hit to a CR or from a CR to a hit may involve a task switch, and definitively requires a response switch (between 1s and 2s to 3s and 4s or vice versa). Thus, the observed memory state dissociations may be unrelated to memory content and may instead reflect changing task demands. We can assess memory content by leveraging the confidence judgments made for each test-phase response. Our assumption is that higher confidence responses reflect greater memory content or evidence^13, 14^. If the observed pattern is solely the result of changing task demands, then memory state evidence should not be modulated by response confidence. Alternatively, memory state engagement may be modulated by memory content such that the congruency effect is greater for either high or low confidence responses.

To test the extent to which memory content interacts with congruency-dependent memory state engagement, we assessed memory state evidence as a function of prior response, current response, and current response confidence. We averaged response-locked retrieval evidence over the 500ms pre-response interval^15, 16^. We conducted a 2*×*2*×*2 rmANOVA with factors of current response (hit, similar CR), prior response (hit, CR), and confidence (high, low) with memory state evidence in the pre-response interval as the dependent variable. We find a significant main effect of current response (*F* _1,26_=11.22, *p*=0.0025, 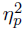=0.3014) driven by greater retrieval evidence for similar CRs (M=0.0778, SD=0.0533) compared to hits (M=0.0454, SD=0.0635). We do not find a significant main effect of prior response (*F* _1,26_=0.0001, *p*=0.9938, 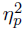 *<*0.0001). We find a significant main effect of confidence (*F* _1,26_=9.003, *p*=0.0059, 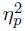=0.2572), whereby retrieval evidence is greater for low confidence responses (M=0.0837, SD=0.0976) compared to high confidence responses (M=0.0395, SD=0.0844). As in the stimulus-locked analysis, we find a significant interaction between current response and prior response (*F* _1,26_=34.98, *p<*0.0001, 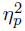=0.5736). We find a significant interaction between current response and confidence (*F* _1,26_=8.794, *p*=0.0064, 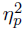=0.2528), but do not find a significant interaction between prior response and confidence (*F* _1,26_=0.7051, *p*=0.4087, 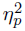=0.0264). We find a significant three way interaction between current response, prior response, and confidence (*F* _1,26_=5.613, *p*=0.0255, 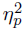=0.1776).

The confidence analysis suggests the existence of a *mnemonic congruency effect* whereby memory content influences mnemonic state engagement selectively for congruent trials. To directly test this account, we conducted paired *t* -tests on retrieval evidence for high and low confidence responses separately for congruent and incongruent trials. For congruent trials, we find greater retrieval evidence for low (M=0.0690, SD=0.0832) compared to high (M=0.0024, SD=0.0509) confidence responses (*t* _26_=-3.891, *p*=0.0006, *d* =0.9654). For incongruent trials, we do not find a significant difference between high (M = 0.0765, SD=0.0629) and low confidence responses (M=0.0984, SD=0.0883; *t* _26_=-1.224, *p*=0.2321, *d* =0.9654). To directly test the specificity of this confidence effect, we computed difference scores (high-low confidence) for congruent and incongruent trials. Paired *t* -tests revealed significantly larger difference scores for congruent (M=-0.0666, SD=0.0509) compared to incongruent (M=-0.0219, SD=0.0912) trials (*t* _26_=-2.369, *p*=0.0255, *d* =0.5009). Together, these results indicate that the effect of confidence is specific to congruent trials.

## Discussion

The aim of the present study was to understand how prior memory responses influence subsequent behavior and memory state dynamics. We directly tested the hypothesis that prior responses induce ‘lingering’ brain states that drive changes in current behavior. We used scalp EEG and an independently validated mnemonic state decoder to measure memory brain states while participants completed a recognition memory task. We replicate past behavioral findings that prior memory responses influence current behavior^7^. However, instead of finding that ‘old’ judgments induce a retrieval state and ‘new’ judgments induce an encoding state, we find that all test-phase judgments induce a retrieval state which does not appear to linger in time. Furthermore, retrieval state engagement on the current trial is strongly driven by *congruency* with respect to the prior trial – when the test probes switch across two consecutive trials, we observe greater retrieval state engagement. Finally, we find that this effect of congruency interacts with response confidence. Together, these results suggest that cortical memory states are engaged with respect to the *intention* to retrieve, and potentially reflect the attentional or control demands needed to switch into and maintain hippocampal memory states.

Replicating past behavioral work^7, 8^, we find that prior responses influence current behavior. Specifically, test probes are more often correctly identified when the prior response and current probe are congruent. A target is more likely to be correctly and confidently identified as “old” when the prior response was a hit, and a similar lure is more likely to be correctly and confidently identified as “new” when the prior response was a correct rejection (CR). We did not replicate the finding that these effects are specific to fast (*<* 500ms) inter-stimulus intervals and instead find this effect for all ISIs. Yet as suggested by prior work, these results imply a lingering influence of prior responses on the current task.

Based on prior behavioral work, which we replicate here, we expected to find that hits would induce a lingering retrieval state and CRs would induce a lingering encoding state, yet our results run counter to these predictions. We find retrieval state engagement for both hits and CRs, with greater retrieval state engagement for CRs, and we find that memory state evidence is near to zero following all memory responses. Our interpretation of these surprising findings is that we are observing the engagement of a brain state recruited to support the *attempt* to retrieve. The initial conceptualization of the retrieval mode is that it constitutes a state engaged in the service of explicit, episodic retrieval^17, 18^. Based on this framing and subsequent work^19–21^, the retrieval state should therefore be observed any time an individual is faced with the top-down demand to retrieve a past experience. Our interpretation is that retrieval state engagement across both hits and CRs in the current study reflects a top-down, intentional goal to retrieve, consistent with our prior work^22^. We observe engagement of the retrieval state for both hits and CRs, which we believe reflects the top-down demand to retrieve old items throughout the test-phase. We additionally find no evidence that the memory states we measure linger in time. We included a variable ISI based on past work which showed that effects of the prior trial diminish with ISIs longer than 500ms; however, neither behaviorally nor neurally do we find interactions with ISI. We speculate that the lack of evidence for lingering memory states may be due to the unpredictable nature of the ISI. Because on any given trial the ISI could be 250ms, participants may always prepare for such an ISI, quickly exiting the retrieval state in anticipation of the onset of the next trial. A shift towards encoding may reflect preparation for the next trial^23^.

Although we do not find evidence for lingering memory states across consecutive trials, we do find that prior responses influence memory state engagement on the current trial, albeit not in the predicted direction. Specifically, we find robust retrieval state engagement on *incongruent* trials, that is, during hits preceded by CRs and CRs preceded by hits. The logic of our initial predictions was based on the notion that the act of recognizing a probe as ‘old’ pushes the hippocampus into a pattern-completing retrieval state^24–26^, which may then persist and facilitate pattern completion when the subsequent probe is presented^7, 8^. It is possible that such a hippocampal mechanism is engaged in this task, but that what we measure at the cortex does not directly reflect these hippocampal memory states. Instead, our results are more consistent with the interpretation that the currently observed cortical brain states reflect the degree to and direction in which *attention* is being deployed. It is well established that cognitive control and/or attentional mechanisms are recruited when an individual must switch between tasks^27–29^. The incongruent trials here may reflect task switching whereby more control or attention is required when the test probe switches across two consecutive trials. The question then becomes, why do attentional demands recruit a putative memory retrieval state? Our interpretation is that when the probe switches, participants must engage internally directed attention^3, 30–32^ in order to initiate the task-relevant memory state. Intriguingly, the cortical brain state that we observe may be a response to conflict created by a lingering, task-irrelevant hippocampal state. That is, the hippocampus may remain in a retrieval or encoding state driven by the prior trial – which would be consistent with the literature and our *a priori* expectations – but when task demands change across trials, an attentional/control state is needed to shift the hippocampus into the task-relevant memory state. A targeted investigation of the link between subcortical and cortical brain states is warranted given these findings.

Across all response types, we find that memory state evidence fluctuates across the stimulus interval. Specifically, we find robust decreases in retrieval state evidence early in the stimulus interval. Due to the nature of our cross-study mnemonic state classifier, this decrease corresponds to an increase in evidence for an encoding state. Early engagement of the encoding state may reflect perceptual processing of and external attention to the test probe^33, 34^. Intriguingly, encoding state evidence appears to dissociate current hits based on prior responses, such that hits preceded by hits recruit the encoding state to a greater degree than hits preceded by CRs. This finding suggests that the reduced demand to switch task sets between consecutive hits may enable attentional orienting to the external world instead, increasing the fidelity of the information that is ultimately sent to the hippocampus to be used in pattern-completion and retrieval. The role of the encoding state, and its link to external attention, is an important avenue for future work.

Finally, we find evidence for a mnemonic congruency effect. We find that although retrieval state evidence is lower for congruent compared to incongruent trials, within congruent trials low confidence responses recruit the retrieval state to a greater degree than high confidence responses. The dissociation in confidence replicates recent work from our lab^22^ and is consistent with the interpretation that the retrieval state is engaged in the service of the attempt to retrieve. To the extent that high confidence responses are supported by greater reinstatement and/or evidence compared to low confidence responses^35, 36^, retrieval state engagement may prolong memory search and/or evidence accumulation processes^14, 37^ that are needed to make a decision. That we observe a dissociation in confidence on congruent trials indicates that retrieval state engagement is not solely driven by task switching. A task switch alone could explain the observed behavioral effects as it is well established that switch trials lead to behavioral costs^28, 38^. That the observed dissociation in confidence appears specific to congruent trials suggests that a single internal attention state may support both task switching and evidence accumulation. We might have anticipated finding the greatest degree of retrieval state evidence for incongruent, low confidence trials, whereby participants engage two processes: one to initiate the task set and one to accumulate evidence. Instead, we do not find a significant confidence-driven difference in retrieval state evidence on incongruent trials. Thus, it may be that a shift between different types of internal information (task set vs. mnemonic representation) does not require a full-scale re-configuration of brain states if internal attention is already engaged.

In summary, we demonstrate that ongoing memory processing is modulated by a combination of current memory demands and prior behavior. Our results suggest that memory brain states may be recruited in response to changes in attentional and memory demands, and support both memory and broader goals. These results advance our understanding of how memory brain states are engaged in response to a dynamic environment.

## Methods

### Participants

27 (22 female, age range = 18-23, mean age = 20.22 years) fluent English speakers from the University of Virginia and surrounding Charlottesville community participated. Our sample size was determined *a priori* based on behavioral pilot data (N=10) described in the pre-registration report of this study (https://osf.io/2v8mn). No participants were excluded from the final dataset. All participants had normal or corrected to normal vision. Informed consent was obtained in accordance with University of Virginia Institutional Review Board for Social and Behavioral Research and participants were compensated for their participation. All raw, de-identified data and the associated experimental and analysis codes used in this study will be made available via the Long Term Memory Lab Website upon publication.

### Experimental Design

#### Recognition Task

In the present study, participants completed a recognition memory task containing a study phase and a test phase. Stimuli consisted of images of common objects drawn from an image database with multiple exemplars per object category^39^.

*Study Phase*: During the study phase, participants studied 16 lists of 18 images of common objects in anticipation of a later memory test, for a total of 288 images (Figure 1A). Each trial consisted of an image of a single, unique object on a white background. Each image was presented for 2000 ms, with a 1000 ms interstimulus interval (ISI). Participants made no responses during the study phase. Participants were asked to look at each image, think about it, and try to remember it in anticipation of a later memory test.

*Test Phase*: Following the final study list, participants completed the test phase, in which the 288 images from the study phase (targets) were presented along with 288 new object images that were not presented during the study phase. Half of the new objects (n=144) were *similar lures*, new images from the same object category as an object presented during the study phase. For example, if a participant saw a bench during the study phase, the similar lure would be an image of a different bench. The other half of the new objects (n=144) were *novel items*, images from an entirely new object category (e.g. a fan, assuming that no fan image was presented during study). In the test phase, participants were presented with a single image and asked to indicate their confidence that the image was ‘old’ (previously studied) or ‘new’ (not previously studied) on a scale of 1-4, whereby 1 indicated a confident ‘new’ judgment, 4 indicated a confident ‘old’ judgment, and 2 and 3 were uncertain ‘new’ and ‘old’ judgments, respectively. During the test phase, the ISI between test probes varied from 250ms to 1000ms in 250ms intervals (250ms, 500ms, 750ms, 1000ms). ISI varied randomly, whereby each ISI was equally likely to occur following targets, similar lures, and novel items.

#### Mnemonic State Task

An independent group of participants completed a mnemonic state task. Participants were biased via explicit instructions on a trial-by-trial basis to engage an encoding or retrieval state, while perceptual input and behavioral demands were held constant. In this mnemonic state task (for specific study parameters, please see^6, 11^), participants viewed two lists of object images. For the first list, each object was new. For the second list, each object was again new but was categorically related to an object from the first list. For example, if List 1 contained an image of a bench, List 2 would contain an image of a different bench. During List 1, subjects were instructed to encode each new object. During List 2, however, each trial contained an instruction to either encode the current object (e.g., the new bench) or to retrieve the corresponding object from List 1 (the old bench). Each object was presented for 2000 ms. Participants completed either a two-alternative forced choice recognition test or a recency test on the object stimuli. We used the stimulus-locked List 2 data, data that is time-locked to when the stimulus appears on the screen, to train a multivariate pattern classifier (see below) to distinguish encoding and retrieval states.

### EEG Data Acquisition and Preprocessing

All acquisition and preprocessing methods are based on our previous work^6^. EEG recordings were collected using a BrainVision system and an ActiCap equipped with 64 Ag/AgCl active electrodes positioned according to the extended 10-20 system. All electrodes were digitized at a sampling rate of 1000 Hz and were referenced to electrode FCz. Offline, electrodes were later converted to an average reference. Impedances of all electrodes were kept below 50kΩ. Electrodes that demonstrated high impedance or poor contact with the scalp were excluded from the average reference. Bad electrodes were determined by voltage thresholding (see below).

Custom Python codes were used to process the EEG data. We applied a high pass filter at 0.1 Hz, followed by a notch filter at 60 Hz and harmonics of 60 Hz to each participant’s raw EEG data. We then performed three preprocessing steps^40^ to identify electrodes with severe artifacts. First, we calculated the mean correlation between each electrode and all other electrodes as electrodes should be moderately correlated with other electrodes due to volume conduction. We z-scored these means across electrodes and rejected electrodes with z-scores less than -3. Second, we calculated the variance for each electrode, as electrodes with very high or low variance across a session are likely dominated by noise or have poor contact with the scalp. We then z-scored variance across electrodes and rejected electrodes with a *|*z*| ≥* 3. Finally, we expect many electrical signals to be autocorrelated, but signals generated by the brain versus noise are likely to have different forms of autocorrelation. Therefore, we calculated the Hurst exponent, a measure of long-range autocorrelation, for each electrode and rejected electrodes with a *|*z*| ≥* 3. Electrodes marked as bad by this procedure were excluded from the average re-reference. We then calculated the average voltage across all remaining electrodes at each time sample and re-referenced the data by subtracting the average voltage from the filtered EEG data. We used wavelet-enhanced independent component analysis^41^ to remove artifacts from eyeblinks and saccades.

### EEG Data Analysis

For the recognition task data, we applied the Morlet wavelet transform (wave number 6) to the entire EEG time series across electrodes, for each of 46 logarithmically spaced frequencies (2-100 Hz;^42^). We then downsampled the test-phase data by taking a moving average across 50 ms time intervals from -1000 to 3000 ms relative to the response and sliding the window every 25 ms, resulting in 314 time intervals (80 non-overlapping). Mean and standard deviation power were calculated across all trials and across time points for each frequency. Power values were then z-transformed by subtracting the mean and dividing by the standard deviation power. We followed the same procedure for the mnemonic state task, with 317 overlapping (80 non-overlapping) time windows from 4000 ms preceding to 4000 ms following stimulus onset^6, 16^.

### Pattern Classification Analysis

We conducted pattern classification analysis using penalized (L2) logistic regression, implemented via the sklearn linear model module in Python and custom Python code. For all classification analyses, classifier features were comprised of spectral power across 63 electrodes and 46 frequencies. Prior to conducting pattern classification analyses, we performed an additional round of z-scoring across features (electrodes and frequencies) to eliminate trial-level differences in spectral power. We used classifier evidence as a trial-specific, continuous measure of memory brain state engagement, to asses how memory judgments modulate and are modulated by memory brain states.

#### Cross Study Memory State Classification

To measure memory state engagement in the recognition memory task, we conducted three stages of classification, using the same methods as in our prior work^3, 22^. First, we conducted within subject leave-one-run out cross validated classification (penalty parameter = 1) on all participants who completed the mnemonic state task (N = 100, see^11^ for details). The classifier was trained to distinguish encode vs. retrieve states based on spectral power averaged across the 2000 ms stimulus interval during List 2 trials. For each participant, we generated true and null classification accuracy values. We permuted condition labels (encode, retrieve) for 1000 iterations to generate a null distribution for each participant. Any participant whose true classification accuracy fell above the 90th percentile of their respective null distribution was selected for further analysis (N = 35). Second, we conducted leave-one-subject out cross-validated classification (penalty parameter = 0.0001) on the selected participants to validate our mnemonic state classifier. We found significantly above chance classification accuracy (M = 60%, SD = 9.5%, t_34_ = 6.1507, p *<* 0.0001), indicating that the cross subject classifier is able to distinguish encoding and retrieval states. Finally, we applied the cross subject mnemonic state classifier to the recognition task data, specifically spectral signals in 50 ms time windows. We extracted classifier evidence, the logit-transformed probability that the classifier assigned a given recognition task trial a label of encoding or retrieval. This approach provides a trial-level estimate of memory state evidence during the recognition task. Due to the structure of our classifier, positive values of memory state evidence reflect greater retrieval state engagement and negative values of memory state evidence reflect greater encoding state engagement.

### Statistical Analyses

We used repeated measures ANOVAs (rmANOVAs) to assess the impact of prior judgment and ISI on current judgment accuracy and current judgment confidence. We used rmANOVAs to assess the impact of both prior and current memory judgments on memory brain states. We followed up significant interactions with paired *t* -tests. We used false discovery rate (FDR) to correct for multiple comparisons for post-hoc *t*-tests^43^. We report effect sizes as partial eta squared (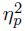) and Cohen’s *d*. We conducted Bayes Factor analysis using the BayesFactor package (version 0.9.12-4.4) in R (version 4.2.3) with the default prior settings and specifically the linear model function lmBF to compare models.

## Acknowledgments

We thank Yuju Hong and Devyn Smith for assistance with data collection. This work was supported by a grant from the National Institutes of Health (NINDS R01 NS132872, PI: NML) and a Harrison Award from the University of Virginia Office of Undergraduate Research.

## Author contributions

Justin R. Wheelock: Conceptualization; Data curation; Formal analysis; Funding acquisition; Software; Visualization; Writing—Original draft; Writing—Review & editing. Nicole M. Long: Conceptualization; Formal analysis; Funding acquisition; Project administration; Software; Supervision; Visualization; Writing—Original draft; Writing—Review & editing.

## Declaration of interests

The authors declare no competing interests.

## Data and code availability

The datasets generated in the current study, along with all experimental codes used for data collection and data analysis will be made available on the Open Science Foundation (OSF) via the Long Term Memory Lab website (http://longtermmemorylab.com/publications/) upon publication.

## Notes

### Competing Interest Statement

The authors have declared no competing interest.

